# Analysis of circulating-microRNA expression in lactating Holstein cows under summer heat stress

**DOI:** 10.1101/2020.03.18.996777

**Authors:** Jihwan Lee, Suhyun Lee, Junkyu Son, Hyeonju Lim, Euntae Kim, Donghyun Kim, Seungmin Ha, Taiyoung Hur, Seunghwan Lee, Inchul Choi

**Affiliations:** Dairy Science Division, National Institute of Animal Science, RDA, Cheon-an, Republic of Korea; Department of Animal and Dairy Sciences, Chungnam National University, Republic of Korea

## Abstract

Korean peninsular weather is rapidly becoming subtropical due to global warming. In summer 2018, South Korea experienced the highest temperatures since the meteorological observations recorded in 1907. Heat stress has a negative effect on Holstein cows, the most popular breed of dairy cattle in South Korea, which is susceptible to heat. To examine physiological changes in dairy cows under heat stress conditions, we analyzed the profiles circulating microRNAs isolated from whole blood samples collected under heat stress and non-heat stress conditions using small RNA sequencing. We compared the expression profiles in lactating cows under heat stress and non-heat stress conditions to understand the regulation of biological processes in heat-stressed cows. Moreover, we measured several heat stress indicators, such as rectal temperature, milk yield, average daily gain, and progesterone concentration. All these assessments showed that pregnant cows were more susceptible to heat stress than non-pregnant cows. Particularly, progesterone concentrations known to have maternal warming effects were at similar levels in non-pregnant cows but significantly increased in pregnant cows under heat stress conditions. The differentially expressed miRNAs and their putative target genes were analyzed in pregnant cows. Interestingly, we found that differentially expressed miRNAs (bta-miR-146b, bta-miR-20b, bta-miR-29d-3p, bta-miR-1246) specifically targeted progesterone biosynthesis (*StAR*) and the function of corpus luteum-related genes (*CCL11*, *XCL*), suggesting that pregnant cows with elevated progesterone concentrations are more susceptible to heat stress. In addition, we found the differential expression of 11 miRNAs (bta-miR-19a, bta-miR-19b, bta-miR-30a-5p, and several from the bta-miR-2284 family) in both pregnant and non-pregnant cows under heat stress conditions. In target gene prediction and gene set enrichment analysis, these miRNAs were found to be associated with the cytoskeleton, cell junction, vasculogenesis, cell proliferation, ATP synthesis, oxidative stress, and immune responses involved in heat response. These miRNAs can be used as potential biomarkers for heat stress.

## Introduction

Following global warming, including the northward expansion of the subtropical climate zone, South Korea, located at the northern hemisphere of East Asia, is susceptible to the impact of climate change [1–3]. For example, the average increase in temperature in South Korea was 1.7°C, while global temperature increased by 0.7°C from 1912 to 2008 [2, 4]. Mainly, in summer 2018, South Korea experienced extreme hot temperatures since the meteorological observations recorded in 1907. Moreover, heat stress (HS) has a negative influence on livestock productivity, particularly, Holstein cows (*Bos Taurus*), the most popular breed of dairy cattle in South Korea. They are more sensitive to HS as it induces hormonal changes, infection, metabolic disorders, and abnormal embryo development in dairy cattle [5, 6], consequently affecting the economic traits such as growth, milk production, and infertility. HS can lead to limited feed-intake and imbalanced hormone secretion, resulting in a decrease in growth and reproductive efficiency. For example, the placental function of heat-stressed cows during late gestation is impaired by decreased secretion of placental hormones such as estrone sulfate, leading to retarded fetal growth and low birth weight of the calves [7]. Moreover, the reproductive performance of Holstein cow is more susceptible to HS during summer, suggesting that HS conditions, including elevated temperature and humidity, decrease thermal tolerance [8]. Jiangijing Liu et al. (2019) reported that the pregnancy rate of Holstein cows is 39.4% at temperature-humidity index (THI) < 72 (non-HS) and decreased to 31.6% at THI > 78.0 (Intermediate HS) [9]. In Florida, pregnancy rates of lactating cows in the summer is low (13.5%) [10]. Furthermore, the number of mounts per estrus also decreased by nearly half in summer, compared to winter [11]. For lactating cows, HS conditions have been reported to reduce milk yield by about 30-40 % [12–14].

In addition to physiological responses, economic losses, including the cost of veterinary care and farm management (fans, sprinklers installation) and involuntary culling can have negative impacts on the dairy industry [15–17]. Therefore, the development of feasible methods and identification of biomarkers are essential for recognizing heat-stressed cows in order to provide individual attention and tailor-made care. To investigate the effects of heat stress on dairy cows, we used profiles of circulating microRNAs isolated from whole blood that was collected in HS and non-HS season. Recent studies have demonstrated that miRNAs are exported to the extracellular environment through microvesicles such as exosomes and circulate in the blood [18, 19] and have shown tremendous potential as non-invasive biomarkers in human cancers [20–22] and pregnancy [23], estrus [24], and aging [17] in dairy cattle. In addition, miRNAs have different expression patterns under environmental and physiological changes such as HS [25, 26]. The goal of this study was to find the potential biomarkers related to HS in lactating dairy cows and to identify the association of candidate miRNAs with putative targeted genes under HS.

## Materials and Methods

### Experimental animals

Before the start of the experiments, veterinarians regularly checked the cows’ medical condition in the dairy research center, and nine lactating Holstein-Friesian cows determined to be healthy and free of disease were selected. All cows had 227±45.5 (mean±standard deviation) average milking days, four of them were pregnant and the others and had a Body Condition Score (BCS) between 3.0 and 3.25. Except for two cows, all cows were similar (S1 Table). Diet was formulated according to NRC 2001, and the cows were fed twice a day to meet the nutrient requirement. Pregnant and non-pregnant cows ate the same feed in the same area. Freshwater was available for free, and mineral blocks were placed on columns of the barn. In this study, all animal experimental designs and procedures were approved by the National Institute of Animal Science Animal Care and Ethics Committee in South Korea (NIAS-109).

### Blood collection

Whole blood was collected from the jugular vein of cows (n=9) at 14:00 in the summer (THI: 86.29) and autumn (THI: 60.87) using PAXgene Blood RNA tube (Qiagen, 762165, California, USA) and vacutainer tube containing sodium heparin (BD, vacutainer®, 367874, Franklin Lakes, NJ, USA). PAXgene Blood RNA tubes were stored at −80°C until miRNA extraction. Heparin tubes were immediately centrifuged at 3,000×g for 10 min at 4°C. Extracted plasma was stored at −80°C until progesterone assay.

### HS indicators

#### Temperature-humidity index (THI)

In order to measure the THI in the barn, a THI measuring device (Testo-174d, 5720500, Germany), which automatically recorded the temperature and relative humidity, was attached to columns in three points in the barn. The measurements were recorded a total of 12 times a day at 2 h intervals. THI was calculated using the following formula [27, 28]: THI = (0.8 × temperature(°C) + [(relative humidity(%)/100) × (temperature(°C) − 14.4)] + 46.4.

#### Body Weight & Milk yield measurement

Bodyweight and milk yield were recorded automatically during the milking by the milking robot (Lely Astronaut milking robot, Netherlands) from June to October. We analyzed the average daily gain (ADG) using bodyweight data. Based on the average milk yield for May, the relative average milk yield for each cow from June to October was calculated.

#### Rectal temperature

Rectal temperature was measured at the same time as the blood collection. In order to measure rectal temperature, the feces of cows were removed. Using rectal thermometer (POLYGREEN Co. Ltd, Germany), the rectal temperature was manually measured at 14:00 under HS (THI: 86.29) and NHS conditions (THI: 60.87). For the accuracy, the measurement was repeated three times per cow, and the rectal thermometer was inserted into the rectum more than 15 cm deep. We also calculated the heat tolerance coefficient (HTC) according to the method described by Road A.O [29]; HTC=100−10(RT(°F)−101).

#### Progesterone assay

Progesterone assay was performed using the collected blood (plasma) under HS and NHS conditions. The plasma P4 concentration was quantified using commercial kits (Siemens 06603261 Immulite Progesterone Kit, USA) and Immulite 1000 Immunoassay System (Siemens DPC Cirrus, USA) according to the manufacturer’s instructions. The inter-assay and intra-assay coefficients of variations were 9.5% and 7.7%, respectively.

### MiRNA extraction and cDNA synthesis

MiRNAs were isolated from whole blood using the PAXgene Blood MicroRNA Kit (Qiagen) according to the manufacturer’s instructions. The miRNA concentrations were determined by using NanoDrop (Optigen NANO Q, South Korea), and cDNA was synthesized using the miScript II RT Kit (Qiagen, 218160, California, USA) following the manufacturer’s instructions and stored at −80°C until use. Realtime-qPCR was performed on 11 differentially expressed (DE) miRNAs based on miRNA-seq results (|FC| ≥ 2, P < 0.05) using miScript SYBR Green PCR Kit (Qiagen 218073, California, USA) according to the manufacturer’s instructions with StepOne Applied Biosystems real-time PCR machine (Applied Biosystems, Foster City, CA). All RT-qPCR reactions were performed in triplicates. Endogenous control was bta-miR-128. All primer information used in this experiment is represented in S2 Table.

### miRNA-seq experiment and statistical analysis

We checked miRNA integrity using an Agilent 2100 Bioanalyzer (Agilent Technologies, Santa Clara, CA, USA) with an RNA integrity number greater than or equal to 7. To construct the library, we performed adapter ligation, reverse transcription, PCR amplification and pooled gel purification using Truseq Small RNA Library Prep Kit (Illumina, San Diego, USA). A flow cell containing millions of unique clusters was added to Illumina Hiseq2000 sequencer. Raw sequencing reads of circulating miRNAs extracted from all samples were pre-processed and analyzed using miRDeep2 software. Adapter trimming was performed to remove the adapter sequences attached to the miRNA during small RNA library construction process using Cutadapt v.1.9.1 and then to increase accuracy, trimmed reads (minimum 18 bp) were collected to form a cluster. The pre-processed and clustered reads were aligned with Mus musculus reference genome, and then those reads were aligned with Mus musculus precursor and matured miRNAs extracted from miRBase v21. To detect known and novel miRNAs and estimate their abundance, we used miRDeep2 software. Raw read data for each miRNA were normalized by the total reads of each sample as standardized to reads per million (RPM, miRNA reads counts/total counts of each sample × 1,000,000). We removed miRNAs with zero RPM values for all samples and added 1 for the RPM values of the filtered miRNAs to facilitate log2 transformation. Filtered data were transformed into log values and normalized using the quantile normalization method. For each miRNA, a fold change was calculated between HS samples and NHS samples. We analyzed DE miRNAs based on |FC ≥ 2| and P < 0.05. All data analysis and visualization for DE miRNAs were performed with R.3.1.2.

### Bioinformatics analysis

Two databases, miRmap (v1.1, mirmap.ezlab.org) and TargetScan (v7.2, targetscan.org), were used to predict the target genes for DE miRNAs [30, 31]. Target genes were selected based on the miRmap Score ≥ 80 (provided by miRmap program) and context++ score percentile ≥ 95 (provided by TargetScan program) was taken as the cut-off value to increase the accuracy of the analysis. Gene Ontology analysis was performed using the PANTHER Classification System (v.14.1) [32] to identify the functional enrichment for a gene set. In addition, we analyzed the Kyoto Encyclopedia of Genes and Genomes (KEGG) pathways by uploading the target gene list to DAVID Bioinformatics Resources 6.8 [33] to identify the functional enrichment signaling pathways related to these target genes.

## Results

### Estimation of THI and blood collection

We measured ambient temperature and relative humidity inside the barn daily to estimate THI (Fig 1). We observed that minimum THI exceeded 72 from the first week of July and even reached over 80, and moderate/severe HS conditions (THI > 78) lasted for over one month. Both maximum and minimum THI peaked around mid-August, and gradually declined; however, mild to moderate HS (THI 72-78) was detected until the end of September. We collected whole blood at 14:00 from both pregnant and non-pregnant cows in the summer (HS) and autumn (NHS). The THI ranged from 79.10 to 87.73 (14:00; 86.29) and 47.3-64.85 (14:00; 60.87) at the HS and NHS sample collecting day. The minimum THI of more than 72 (the cut-off level for HS) until the HS sampling lasted for 36 days and the maximum THI of less than 72 until NHS sampling lasted for 28 days.

**Fig 1.**
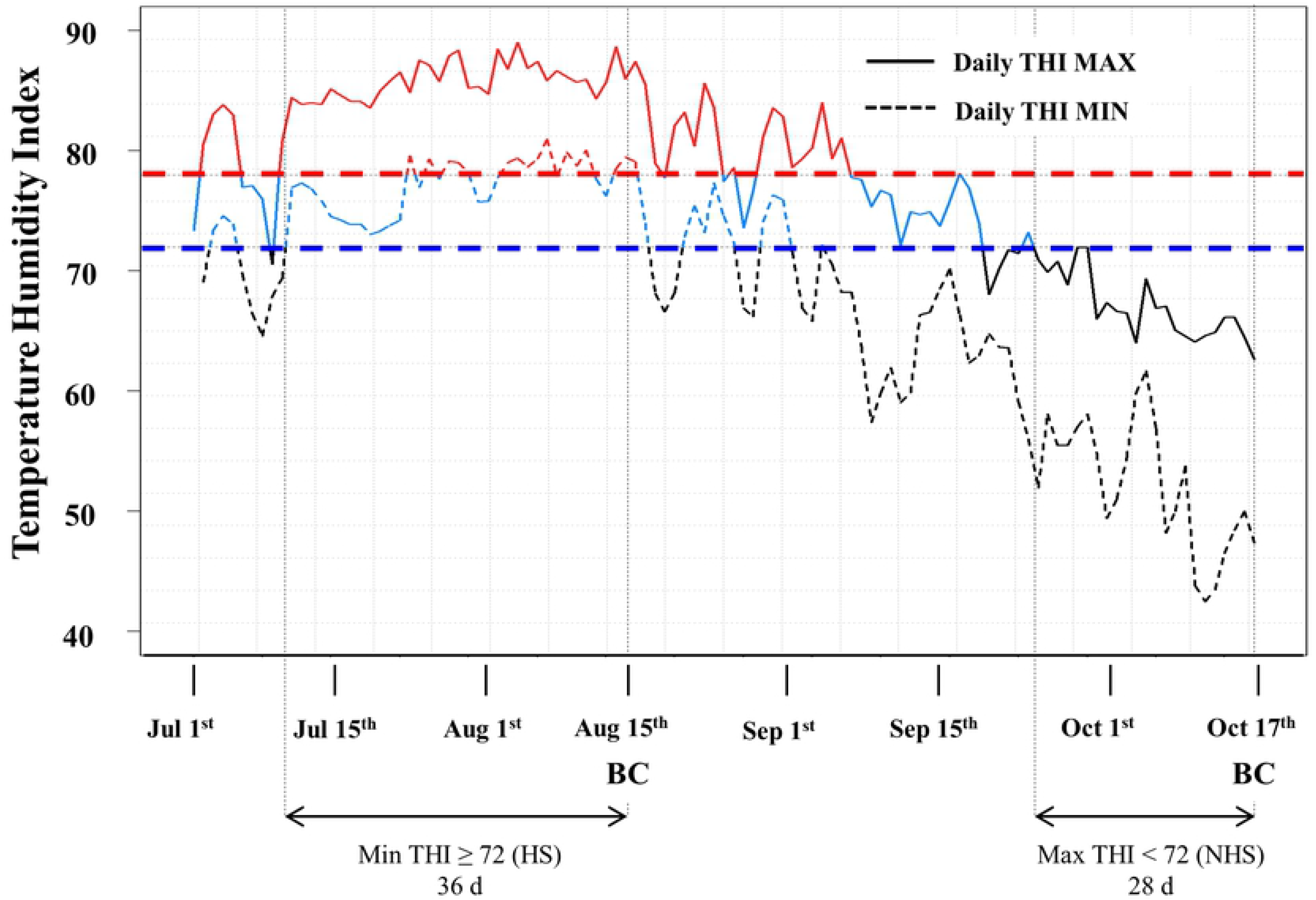
Temperature-Humidity index (THI) measured on dairy barn. The straight line represents daily THI maximum and the dotted line represents daily THI minimum. THI over 78 is marked in red (moderate to severe stress), and 72-78 in blue (mild to moderate stress), and less than 72 in black (non-stress). Vertical dotted line represents THI on the day of sampling. BC, Blood collection; HS, Heat stress condition; NHS, Non-heat stress condition.

### Effects of HS on physiological changes

We measured physiological HS indicators such as rectal temperature, milk yield, ADG of weight, and progesterone concentrations in pregnant and non-pregnant cows. The rectal temperature of cows was measured at the time of blood collection. In NHS conditions, there were no differences between pregnant (38.4°C±0.07) and non-pregnant cows (38.02°C±0.14). However, the rectal temperatures of pregnant cows (40.15°C±0.17) were higher than that of non-pregnant cows (39.36°C±0.1) (P < 0.05) (Table 1). We also observed similar results in the HTC test; no significant differences under NHS conditions, but higher tolerance in non-pregnant under HS conditions were observed (Table 1).

**Table 1.** Rectal temperature and HTC values under environmental conditions on the day of sampling.

|  | Group | Rectal temp. (°C) | HTC | Temperature Range (°C) | THI |
| --- | --- | --- | --- | --- | --- |
| HS | Pregnancy | 40.15±0.17 | 67.3±0.3 | 27.3 - 37.4 | 79.10 - 87.73 |
|  | Non-pregnancy | 39.36±0.11 | 81.52±0.20 |  |  |
| NHS | Pregnancy | 38.40±0.07 | 98.8±0.12 | 8.4 - 17.6 | 47.3 - 64.85 |
|  | Non-pregnancy | 38.02±0.14 | 105.64±0.25 |  |  |

HS, Heat stress; NHS, Non-heat stress; HTC, Heat tolerance coefficient: $100-10 \times (RT(^{\circ}F)-101)$ .

To estimate the effect of HS on milk production, we used relative average milk yield; the ratio of the total milk yield in a month of the testing period to the yield of a month (May). Relative average milk yield decreased gradually in both pregnant and non-pregnant cows, but there was a significant reduction in pregnant cows during extremely hot summer, and the milk production increased after HS (Fig 2A). To investigate the effects of HS on growth performance, we analyzed the changes in ADG of pregnant and non-pregnant cows during testing periods. As shown in Fig 2B, we found an increase in ADG in both pregnant and non-pregnant cows, except for HS pregnant cows during July, although ADG during summer was significantly reduced. We also examined progesterone concentration and found higher concentrations in heat-stressed pregnant cows (6.24±0.362), compared to non-heat-stressed pregnant ones (5.59±0.24).

**Fig 2.**
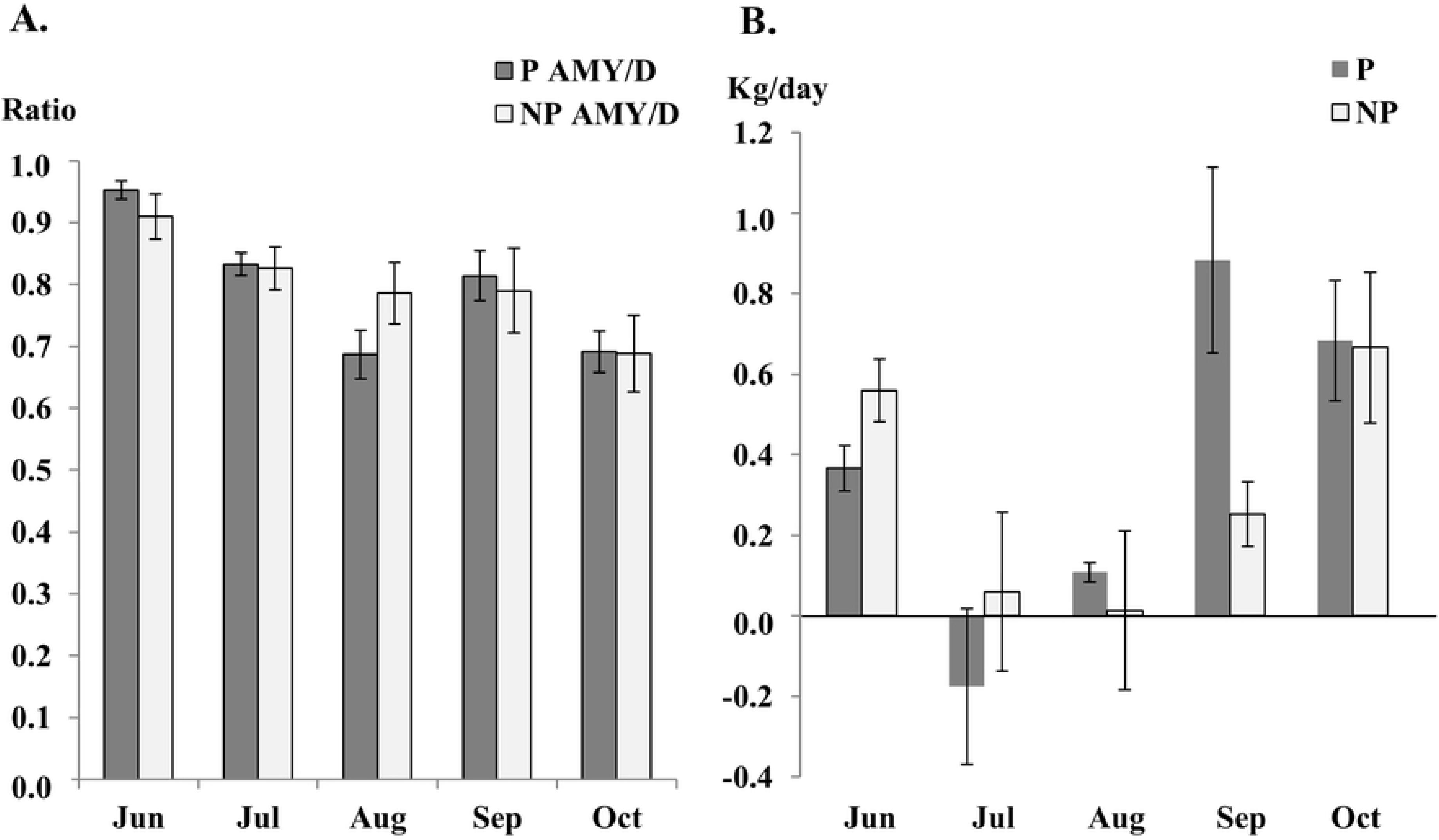
Physiological heat stress indicators recorded during HS and NHS conditions in both pregnant and non-pregnant lactating cows. (A) Relative average milk yield (AMY) to May; (B) Average dairy gain (ADG). HS, Heat stress; NHS, Non-heat stress; P, Pregnancy; NP, Non-pregnancy

### Identification of DE miRNAs of HS cows

We analyzed miRNAs isolated from the whole blood to identify DE miRNAs between heat-stressed and non-heat-stressed cows using small RNA sequencing. In the non-pregnant group, we found that 23 miRNAs were significantly DE (≥ 2-FC in the expression compared to NHS controls; P < 0.05), including nine, upregulated and 14 downregulated miRNAs (Table 2). In the pregnant group, we detected 28 DE miRNAs (10 upregulated, 18 downregulated; ≥ 2-FC and P < 0.05, Fig 3). We identified 11 common DE miRNAs in both non-pregnant and pregnant cows; two miRNAs (bta-miR-19a and bta-miR-19b) were upregulated and nine, including bta-miR-30a-5p and several bta-miR-2284 families, were downregulated (Table 3). We also validated these DE miRNAs by qRT-PCR. The results were very similar to the small RNA sequencing results except for bta-miR-2284x (Table 3).

**Table 2.**
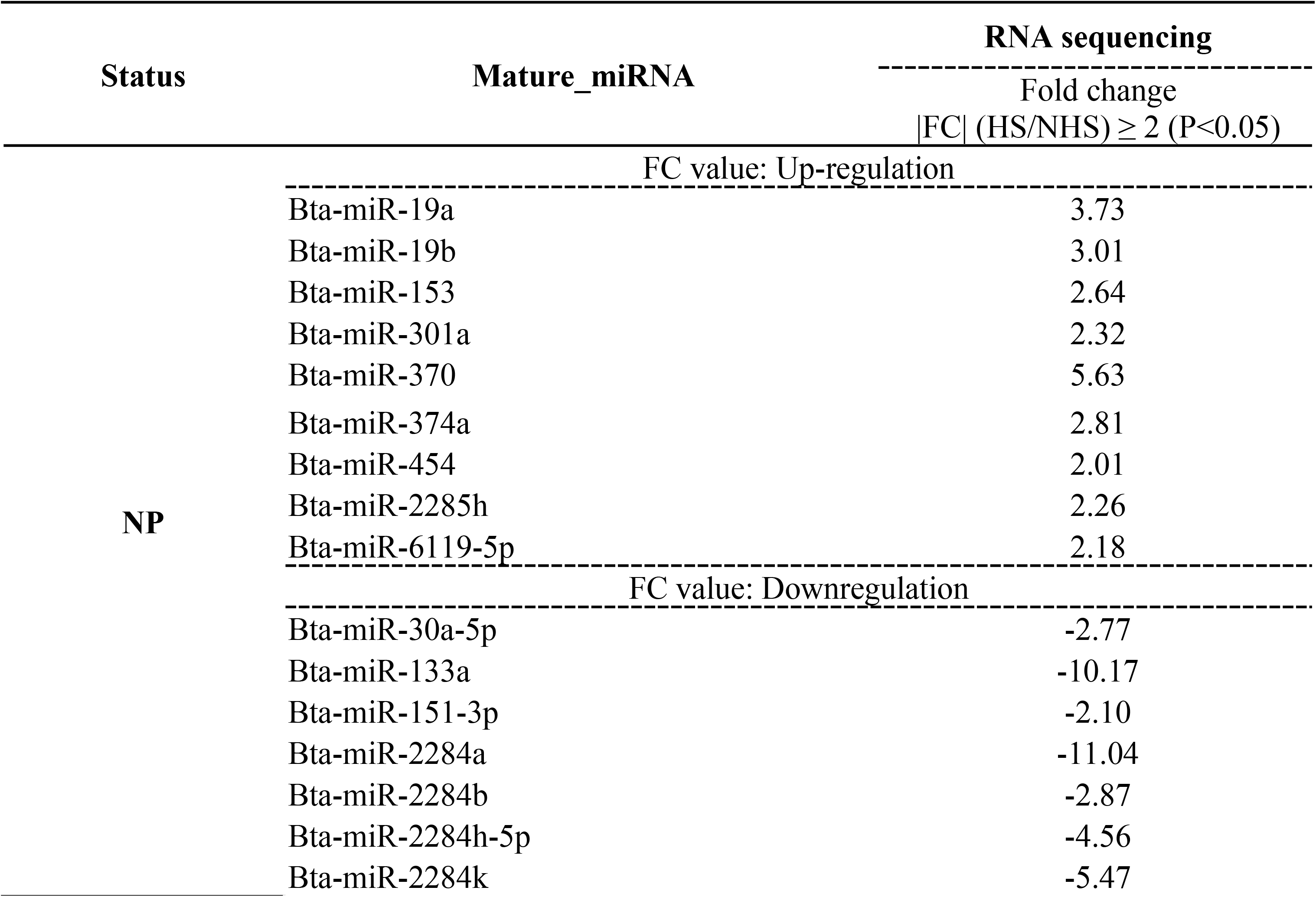
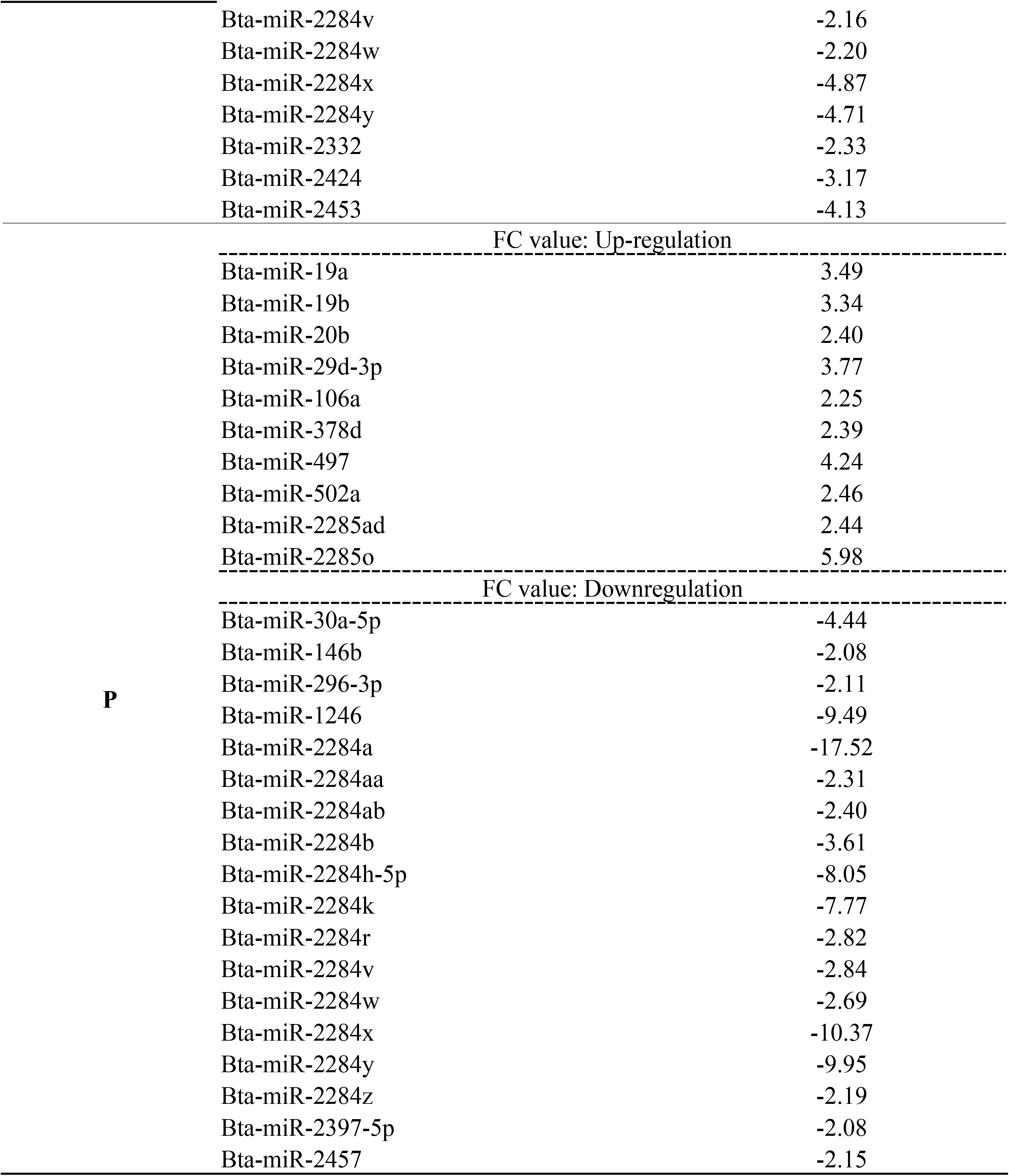
Differentially expressed miRNAs in pregnant and non-pregnant groups under heat-stressed condition.

**Table 3.** Differentially expressed miRNAs in both non-pregnant and pregnant cows under HS conditions (P < 0.05).

| Mature_miRNA | RNA sequencing | RT-qPCR |
| --- | --- | --- |
| | Fold change<br>\|FC\| (HS/NHS) $\geq 2$ (P<0.05) | Endogenous Control: bta-miR-128 |
| **bta-miR-19a** | 3.61 | 2.01 |
| **bta-miR-19b** | 3.17 | 2.14 |
| **bta-miR-30a-5p** | -3.61 | 0.75 |
| **bta-miR-2284a** | -14.28 | 0.42 |
| **bta-miR-2284b** | -3.24 | 0.27 |
| **bta-miR-2284h-5p** | -6.30 | 0.84 |
| **bta-miR-2284k** | -6.62 | 0.98 |
| **bta-miR-2284v** | -2.50 | 0.35 |
| **bta-miR-2284w** | -2.45 | 0.63 |
| **bta-miR-2284x** | -7.62 | 1.08 |
| **bta-miR-2284y** | -7.33 | 0.79 |

HS, heat stress; NHS, non-heat stress

**Fig 3.**
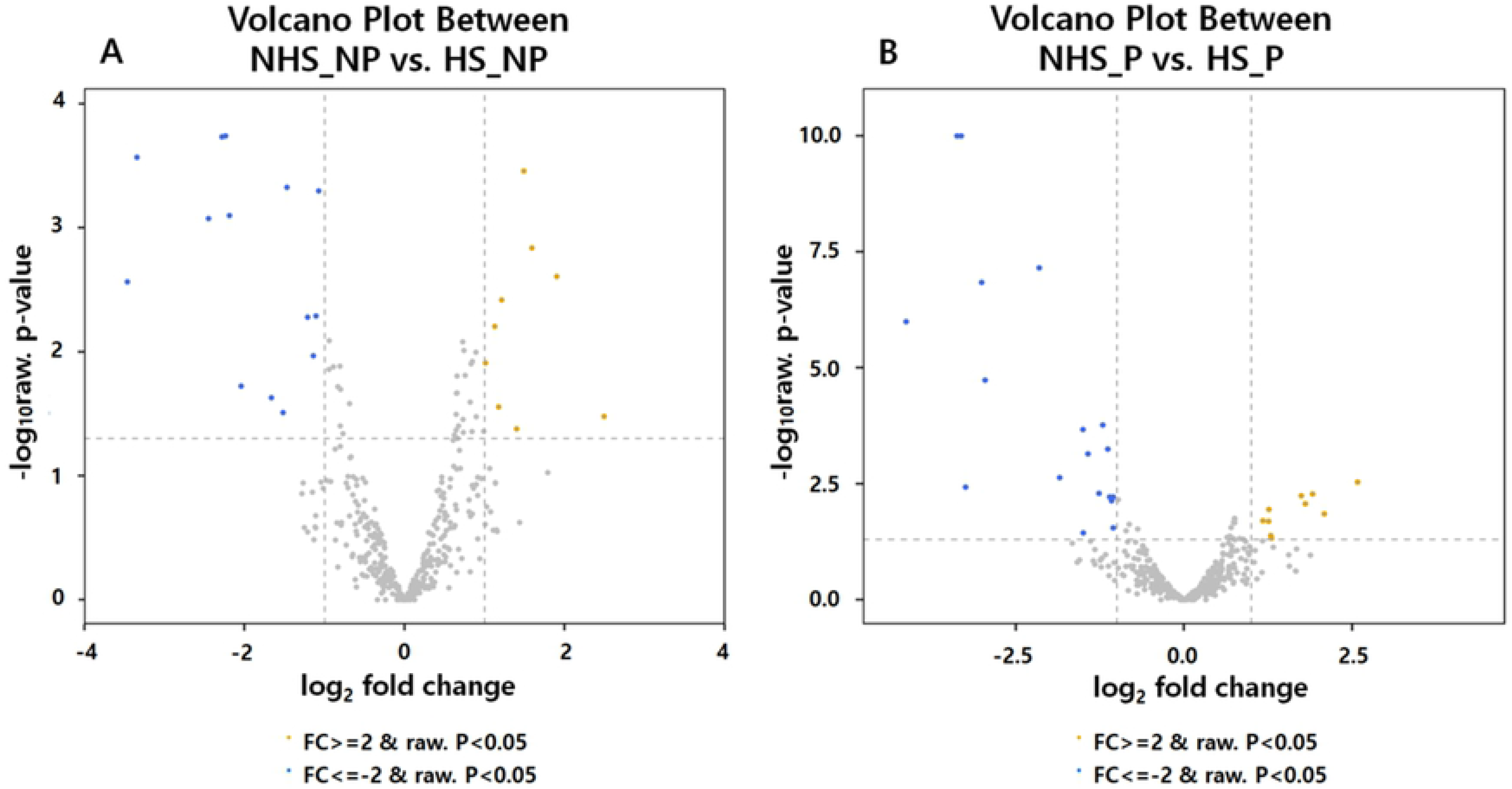
Volcano plot showing differentially expressed miRNAs between NHS and HS using transformed normalized data. |Fold change| value ≥ 2 and P < 0.05 are represented different colors (minus: blue, plus: yellow). (A) Differentially expressed miRNA values in pregnant cows; (B) Differentially expressed miRNA values in non-pregnant cows; HS, Heat stress; NHS, Non-heat stress; P, Pregnancy; NP, Non-pregnancy.

### Putative target gene and signaling pathway analysis

We analyzed putative target genes of 11 common DE miRNAs using miRmap and TargetScan, identified 890 genes, and performed gene set enrichment analysis (GSEA) using DAVID and PANTHER (S1 Fig). The GSEA showed that the putative target genes were associated with the cytoskeleton, cell junction, immune response, oxidative stress involved in heat response (Table 4). Besides, the KEGG pathway showed 18 statistically significant pathways (S3 S1Table). For example, the FoxO signaling pathway, peroxisome, regulation of actin cytoskeleton, TNF signaling pathway, Rap1 signaling pathway, and chemokine signaling pathway were found to be closely related to HS response. Furthermore, we analyzed putative target genes of pregnancy-specific 23 DE miRNAs (S2 Fig). Several DE miRNAs (bta-miR-146b, bta-miR-20b, bta-miR-29d-3p, bta-miR-1246) were found to play essential roles in the regulation of progesterone biosynthesis and corpus luteum (Table 5). KEGG pathway showed statistically significant 24 pathways (Table S4). Interestingly, the prolactin signaling pathway was found to be closely related to progesterone synthesis.

**Table 4.**
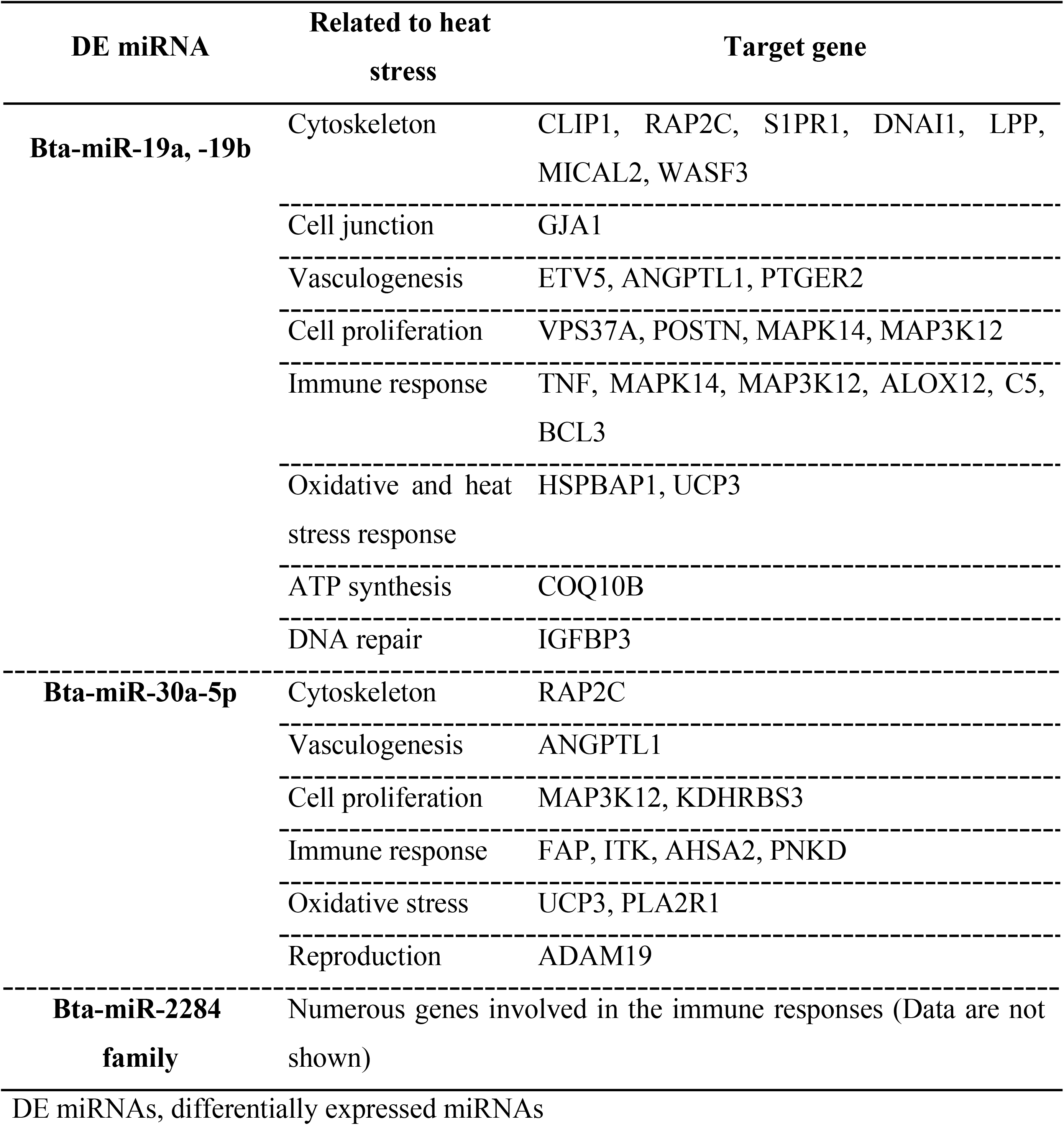
Predicted target genes related to heat stress responses of miRNAs differentially expressed both pregnant and non-pregnant cows.

**Table 5.**
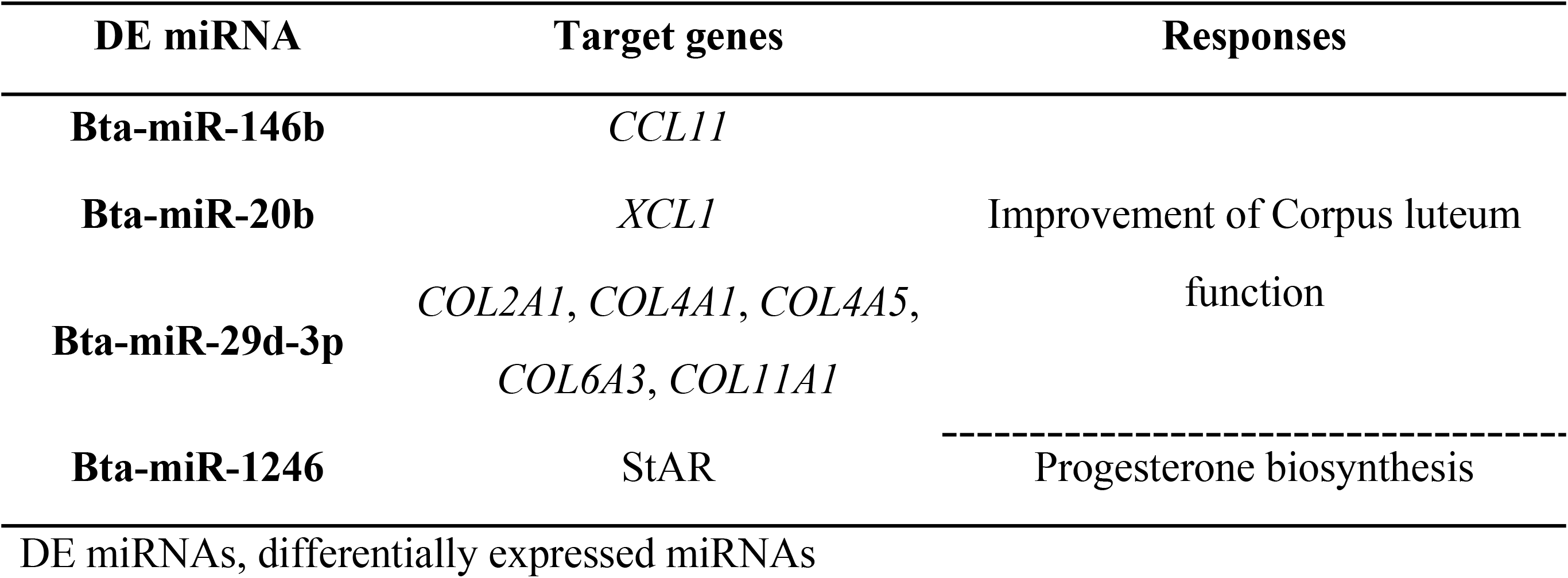
Predicted target genes related to heat stress responses of miRNAs differentially expressed in pregnant cows.

## Discussion

THI is widely used as an indicator to estimate the degree of HS in livestock animals because the valid and reliable assessment of heat load may mitigate or minimize economic loss such as inefficient reproductive performance and milk production in dairy cattle [34, 35]. The THI values can be catabolized into five different classes; no HS (THI < 72), mild HS (72 ≤ THI ≤ 78), moderate HS (78 < THI < 89), severe HS (89 ≤ THI ≤ 98), and death (THI > 98) [36, 37]. To confirm whether cows experienced HS, we checked the association between THI and physiological changes such as milk yield and ADG. We collected whole blood when mild HS lasted more than one month (precisely 36 days), where milk yield and ADG decreased, and non-HS samples were obtained at four weeks after minimum THI returned to < 72 (non-HS condition), implying heat-stressed cows may fully recover from the long-term HS as seen in increased milk production and ADG.

Interestingly, a significant decrease in milk production of pregnant cows during summer might be attributed, in part, to feed-intake reduction because the lactating and pregnant cows need more energy not only for the milk production but also for fetal growth, compared to non-pregnant lactating cows [38, 39]. In this study, we indirectly estimated the feed-intake using ADG. Notably, the negative ADG observed in pregnant HS cows suggested that HS may cause appetite suppression and/or lower feed-intake, consequently resulting in the loss of body weight. The dramatic increase in ADG may be attributed to compensatory placental growth by increased feed-intake.

In accordance with previous studies, where a positive correlation between the rectal temperature and THI value was reported [27], we also observed higher rectal temperature and lower HTC in pregnant HS cows, indicating pregnant cows were more sensitive to HS than non-pregnant cows. In addition, a higher concentration of progesterone in the pregnant HS cows supported our findings because there is a positive correlation between progesterone concentration and rectal temperature during pregnancy [40]. Progesterone secreted from corpus luteum/placenta during pregnancy, and stress-responded progesterone from adrenal glands may be responsible for the increased level [41, 42]. Surprisingly, however, we did not find downregulation of progesterone levels caused by the dysfunction of the corpus luteum during long-term HS. Suggesting the HS cows maintained pregnancy and mild to moderate HS did not impact corpus luteum function and placental development [8, 11]. Physiological indicators proved that pregnant cows were more susceptible to thermal stress, compared to non-pregnant cows.

To further understand the association between these physiological indicators, biological processes, and cellular responses to HS, we identified DE miRNAs using RNAseq. We employed an in silico approach for miRNA target prediction because circulating miRNAs in body fluid may play essential roles in all biological processes and present as potentially useful biomarkers of the HS response. We analyzed 11 common DE miRNAs in both pregnant and non-pregnant cows under HS conditions (Table 3). Two upregulated bta-miR-19a and 19b have been previously reported to target *HSPBAP1*, *DNAJB1*, and *HPX* that respond to heat stress, and downregulated bta-miR-30a-5p may have potential roles in heat stress response, such as oxidation, through its target genes, *PLA2R1* and *PICEN* [25]. In target gene prediction and GESA, bta-miR-19a and bta-miR-19b are associated with biological functions including the cytoskeleton, cell junction, vasculogenesis, cell proliferation, oxidative stress, immune response and ATP synthesis (Table 4). It is well-established that mammalian cells, such as the oocyte, embryo, and mammary gland epithelium were subjected to heat shock. Mainly, degradation and dysfunction of the cytoskeleton in response to HS may result in the aberrant mitochondrial distribution, impaired mitochondrial function, and apoptosis/necrosis. Cellular junctions are also involved in the transportation of ions and small molecules between the blood and milk barrier at the mammary gland. Interestingly, we identified that bta-miR-30a-5p regulated a disintegrin and metalloprotease 19 (*ADAM19*), which is required for early embryo development and implantation in mammals [43]. A ruminant specific miRNA, bta-miR-2284 family, was also identified, putative target genes of which are related to the heat-induced immune response, although the biological roles have not been determined [44].

FoxO signaling pathway (bta04068) has been reported to be associated with several physiological processes, such as oxidative stress response and apoptosis [45]. In HS cells, mitochondria produce large amounts of superoxide radicals, which cause oxidative stress. Oxidative-stressed cells express the molecular chaperone family that are heat shock proteins to protect themselves from protein toxicity. FoxO transcription factor activates the transcription of heat shock proteins such as HSP70, which may serve to protect the cell from protein misfolding associated with oxidative stress [46, 47]. Regulation of actin cytoskeleton pathway (bta04810) is vital in reproduction and milk production of HS cows. In mammalian cells, the cytoskeleton, which is susceptible to heat shock, can degrade or dysfunction under HS conditions, resulting in changes in mitochondria distribution, leading to reduced ATP production [48–50]. Rap1 signaling pathway (bta04015) was found to be closely associated with energy balance control, including glucose homeostasis and lipid metabolism, and it has been reported to play an important role in regulating energy homeostasis in HS conditions [51, 52]. In a previous study, Kim et al. inferred that genes involved in these pathways induce energy homeostasis in response to relatively high energy demands under HS conditions [51]. We also found rapamycin (mTOR) signaling pathway (bta04150), which plays a vital role in several cellular functions such as glucose and lipid metabolism(homeostasis), cellular growth, proliferation, survival, aging, and actin cytoskeletal regulation [53]. Moreover, chemokine signaling pathway (bta04062) and TNF signaling pathway (bta04668), which are associated with inflammatory immune responses, were identified [54, 55]. In our experiment, these signaling pathways were shown to be responsible for several important regulations in lactating cows under HS conditions.

Interestingly, we found that several DE miRNAs (bta-miR-146b, bta-miR-20b, bta-miR-29d-3p, bta-miR-1246) in pregnant cows specifically targeted progesterone biosynthesis (StAR) [56] and function of corpus luteum (*CCL11*, *XCL*, several collagen type genes) related genes (Table 5) [57]. One previous study has reported that when animals are stressed, the adrenal glands that synthesize steroid hormones are stimulated, and the expression levels of *StAR*, which plays an important role in the transformation of cholesterol into progesterone or cortisol, is increased [56]. Moreover, several target genes of pregnancy-specific DE miRNAs have shown to be associated with prolactin signaling pathways (Table S4). Prolactin not only stimulates the secretion of progesterone in corpus luteum but also provides raw material for the synthesis of progesterone [58, 59]. These previous reports support our findings that P4 concentrations increase in summer heat-stressed pregnant cows.

Comprehensively, our findings suggest that the experimentally verified miRNA targets and in silico analysis reflect that the identified miRNA are reliable potential biomarkers of HS response.

## Conclusions

We analyzed several HS indicators such as milk yield, rectal temperature, ADG, progesterone concentration in pregnant and non-pregnant cows and found that pregnant cows are more vulnerable under HS conditions. The DE miRNAs (|FC| ≥ 2, P < 0.05) were analyzed in pregnant cows. Bta-miR-146b, bta-miR-20b, bta-miR-29d-3p, bta-miR-1246 were found to be targeting the genes associated with progesterone biosynthesis and corpus luteum function. This may be related to the increase of progesterone concentration in pregnant cows under HS conditions. Therefore, nutritional strategies and installation of cooling systems (fan, mist sprayer, sprinkler) for managing the heat-stressed pregnant cows may be more important to prevent damage to economic income in the dairy industry. In addition, we also found 11 miRNAs (bta-miR-19a, bta-miR-19b, bta-miR-30a-5p, several from bta-miR-2284 family) that were DE in both pregnant and non-pregnant cows. These 11 miRNAs could be potential biomarkers associated with HS. Moreover, if a large amount of genomic information is collected, genomic selection of heat-tolerant animals and precision feeding management under HS conditions will be possible.

## Acknowledgments

We thank the Dairy Science Division at National Institute of Animal Science for helping us work on this project and JH Kim, a researcher at Macrogen, NGS company located in Daejeon, South Korea for helping us with miRNA-sequencing analysis.

## Author Contributions

Conceptualization: Jihwan Lee, Inchul Choi

Data Curation: Jihwan Lee, Inchul Choi

Formal Analysis: Seunghwan Lee, Suhyun Lee

Funding acquisition: Jihwan Lee

Investigation: Jihwan Lee

Methodology: Jihwan Lee, Inchul Choi

Project administration: Jihwan Lee

Resources: Jihwan Lee, Inchul Choi, Junkyu Son, Hyeonju Lim, Euntae Kim, Donghyun Kim, Taiyoung Hur, Seungmin Ha

Software: Jihwan Lee, Inchul Choi, Seunghwan Lee, Suhyun Lee

Supervision: Inchul Choi

Validation: Jihwan Lee, Inchul Choi

Visualization: Jihwan Lee, Inchul Choi, Seunghwan Lee, Suhyun Lee

Writing-original draft: Jihwan Lee, Inchul Choi

Writing-review&editing: Jihwan Lee, Inchul Choi

## Supporting information captions

**S1 Fig. GO functional analysis for predicted target genes of DE miRNAs in both pregnant and non-pregnant cows.** X axis: Gene ontology classification. Y axis: number of genes in each term.

**S2 Fig. GO functional analysis for predicted target genes of DE miRNAs in pregnant cows.** X axis: Gene ontology classification. Y axis: number of genes in each term.

**S1 Table. Information on each Holstein cows (n=9) used in this experiment**

**S2 Table. Primer Information used for RT-qPCR**

**S3 Table. KEGG pathways enriched for targets of DE miRNAs in both pregnant and non-pregnant cows**. KEGG, Kyoto Encyclopedia of Genes and Genomes.

**S4 Table. KEGG pathways enriched for targets of DE miRNAs in pregnant cows.** KEGG, Kyoto Encyclopedia of Genes and Genomes.

